# Transfer learning via multi-scale convolutional neural layers for human-virus protein-protein interaction prediction

**DOI:** 10.1101/2021.02.16.431420

**Authors:** Xiaodi Yang, Shiping Yang, Xianyi Lian, Stefan Wuchty, Ziding Zhang

**Affiliations:** State Key Laboratory of Agrobiotechnology, College of Biological Sciences, China Agricultural University, Beijing 100193, China; State Key Laboratory of Plant Physiology and Biochemistry, College of Biological Sciences, China Agricultural University, Beijing 100193, China; Dept. of Computer Science, University of Miami, Miami, FL 33146, USA; Dept. of Biology, University of Miami, Miami, FL 33146, USA; Sylvester Comprehensive Cancer Center, University of Miami, Miami, FL 33136, USA

**Author notes:** Corresponding authors (Stefan Wuchty,; Ziding Zhang,;). Other authors (Xiaodi Yang,; Shiping Yang,; Xianyi Lian,).

**Keywords:** human-virus relationship, protein-protein interaction, SARS-CoV-2, prediction, deep learning, transfer learning

## Abstract

To predict interactions between human and viral proteins, we combine evolutionary sequence profile features with a Siamese convolutional neural network (CNN) architecture and a multi-layer perceptron (MLP). Our architecture outperforms various feature encodings-based machine learning and state-of-the-art prediction methods. As our main contribution, we introduce two types of transfer learning methods (i.e., ‘frozen’ type and ‘fine-tuning’ type) that reliably predict interactions in a target human-virus domain based on training in a source human-virus domain, by retraining CNN layers. Our transfer learning strategies can effectively apply prior knowledge transfer from large source dataset/task to small target dataset/task to improve prediction performance. Finally, we utilize the ‘frozen’ type of transfer learning to predict human-SARS-CoV-2 PPIs, indicating that our predictions are topologically and functionally similar to experimentally known interactions. Source code and datasets are available at https://github.com/XiaodiYangCAU/TransPPI/.

## Introduction

Viruses often employ a complex network of protein-protein interactions (PPIs) to coopt their own cellular biological processes, strongly implying that the detection of virus-host PPIs is essential for our understanding of the mechanisms that allow the virus to control cellular functions of the human host. Considerable experimental efforts have been invested in the determination of binary interactions between viral and human proteins, such as yeast two-hybrid (Y2H) assays and mass spectroscopy (MS) technique (Calderwood et al., 2007; Dolan et al., 2013; Gordon et al., 2020; Shah et al., 2019). However, maps of interactions between the human host and various viruses remain incomplete, as a consequence of experimental cost, noise and a multitude of potential protein interactions. Although tens of thousands of interactions have been determined through massive high- and low-throughput experimental technologies, an immense need still exists for the development of reliable computational methods to predict interactions between human and viral proteins.

The primary amino acid sequence remains the most easily to access and complete type of protein information. As a consequence, many sequence-based feature extraction methods have been developed, such as Local Descriptors (LD) (Davies et al., 2008; Yang et al., 2010), Conjoint Triads (CT) (Shen et al., 2007; Sun et al., 2017) and Autocovariance (AC) (Guo et al., 2008; You et al., 2013). Specifically, such features generally represent physicochemical properties or positional information of amino acids that appear in the protein sequences. Based on these feature characterization, several traditional machine learning algorithms (Cui et al., 2012; Dyer et al., 2011; Eid et al., 2016; Emamjomeh et al., 2014; Lian et al., 2020) were previously applied to predict interactions between human and viral proteins. While such machine learning methods effectively captured amino acid specific information to predict novel human-virus PPIs, large data sets impaired their applicability through memory and runtime limits.

In the past decade, applications of deep learning methods have demonstrated improved performance and potential in many fields (e.g., biomedicine, image, and speech recognition) (Karimi et al., 2019; Pospisil et al., 2018; Sainath et al., 2015). In particular, convolutional neural networks (CNNs) (Hashemifar et al., 2018) and recurrent neural networks (RNNs) (Zhang et al., 2016) are comparatively well-established approaches, where CNNs automatically capture local features while RNNs preserve contextualized/long-term ordering information. While deep learning methods (Ahmed et al., 2018; Chen et al., 2019; Du et al., 2017; Hashemifar et al., 2018; Li et al., 2018; Sun et al., 2017) have been designed to predict PPIs with excellent performance such models usually focus on intraspecies interactions. In general, traditional machine learning/deep learning only perform well, if training and test set are cut from the same statistical distribution in the feature space (Shao et al., 2015). While the rigid application of a trained model on testing data sets with different distributions usually perform poorly, transfer learning methods utilize prior knowledge from a ‘source’ to train in a ‘target’ task domain (Chang et al., 2018; Shao et al., 2015) to improve performance. Effective transfer learning can improve the generalization of models, reduce the amount of labeled datasets, and save training time on the target dataset/task. In particular, transfer learning approaches have been successfully applied to problems in many fields, such as medical imaging (Cheplygina et al., 2019), biomedicine (Taroni et al., 2019), and visual categorization (Shao et al., 2015). With the development of deep learning networks, a regular phenomenon appears in various training objectives (Le et al., 2011; Lee et al., 2009) in that the first layers of deep neutral networks (DNNs) usually capture standard features of training data, providing a foundation for transfer learning. Specifically, a deep neural network can be trained on a source task, establishing the parameters of the first layers. Subsequently, parameters of late layers are trained on the target task, striking a balance between the distributions of the different training domains. Depending on the size of the target dataset and number of parameters of the DNN, first layers of the target DNN can either remain unchanged during training on the new dataset or fine-tuned towards the new task, indicating a balance between specificity and generality of derived prior knowledge (Taroni et al., 2019).

Here, we focus on the application of transfer deep learning approaches to the prediction of interactions between proteins of viruses and the human host, an important issue amidst the world-wide COVID-19 pandemic. In particular, we design a deep learning approach to predict interactions between proteins of various viruses and the human host through representing interacting protein sequences with a pre-acquired protein sequence profile module. In particular, we utilize Position Specific Scoring Matrix (PSSM) features to encode the sequence characteristics with a Siamese-based CNN that is fed to a multi-layer perceptron (MLP) (**Figure 1**) (see Materials and methods for details). We propose two types of transfer learning methods through freezing/fine-tuning the parameters of the CNN layers trained with a source and retrained with a target human-virus system to improve prediction performance. Note that we trained the MLP layers with a target human-virus system through randomly initializing parameters. Notably, we found that the transfer of prior knowledge learned from a large-scale human-virus PPI dataset to predict interactions in smaller data sets such as Dengue virus, Zika virus and severe acute respiratory syndrome coronavirus 2 (SARS-CoV-2) improved prediction performance, provided better model generalization and reduced model training time. Finally, we employ the ‘frozen’ transfer learning models to predict human-SARS-CoV-2 PPIs based on pre-trained models with all known human-viral protein interactions. Analysis of the obtained interaction network indicates high functional similarity to experimentally observed host-virus interactions, suggesting that our approach indeed provides meaningful predictions and can guide efforts to understand viral infection mechanisms and host immune responses post SARS-CoV-2 infection.

**Figure 1.**
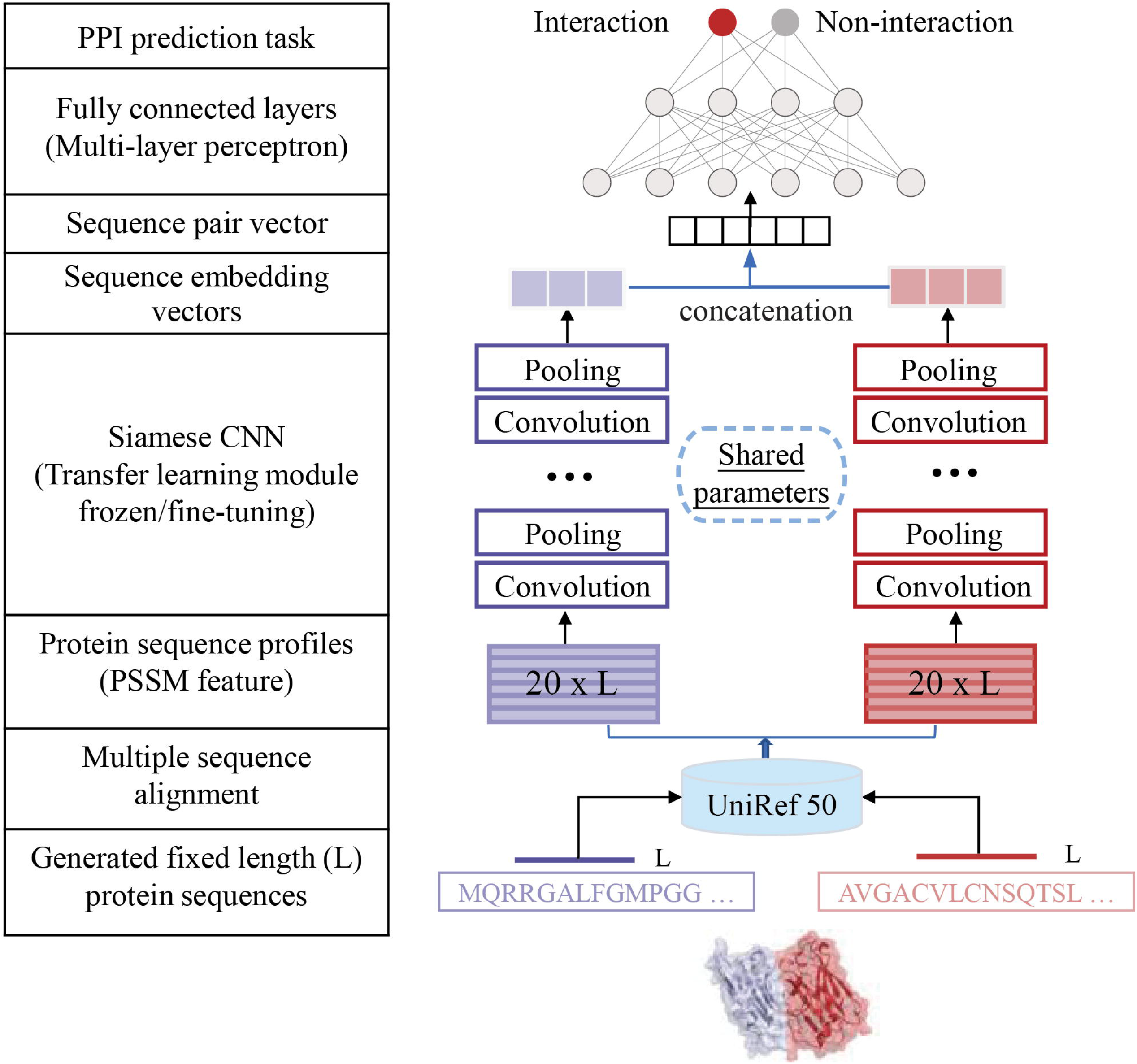
Introducing a deep learning architecture to predict interactions between viral and human proteins, we designed an end-to-end deep neural network framework. In particular, we designed a Siamese CNN module as high-dimensional sequence embeddings that are fed to the prediction module. In particular, the Siamese architecture of the CNN module allows us to account for residual relationships between interacting viral and human protein sequences through position specific scoring matrices (PSSM), capturing evolutionary relationships between proteins. Such latent protein profile representations of interacting protein sequence pairs are fed to the Siamese CNN module to generate respective high-dimensional sequence embeddings. Finally, output embeddings of two proteins are combined to form a sequence pair vector as the input to a multilayer perceptron (MLP) with an appropriate loss function to predict the presence/absence of an interaction between a viral and a human protein.

## Results

### The deep learning framework for human-virus PPI prediction

Representing interactions between human and viral proteins through their sequences, we introduce an end-to-end deep neural network framework, called a Siamese-based CNN that consists of a pre-acquired protein sequence profile module, a Siamese CNN module and a prediction module (**Figure 1**). First, we represent the amino acid composition of interacting proteins by protein sequence profile (i.e., PSSM) features that capture evolutionary relationships. Such latent protein profile representations of interacting protein pairs are fed to the Siamese CNN module to generate respective high-dimensional sequence embeddings. In particular, our CNN module allows us to capture local features of human and viral proteins such as protein linear binding motif patterns, which play important roles in protein interactions. Finally, output embeddings of two proteins are combined to form a sequence pair vector as the input to an MLP with appropriate loss function to predict the presence/absence of an interaction between a viral and a human protein. Details about each module of the deep neural network framework and the implementation details can be found in Materials and methods.

### Performance of the proposed deep learning method

Based on our deep learning architecture we predicted interactions between proteins of various viruses and the human host using 5-fold cross validation. In particular, we utilized 9,880 interactions in HIV, 5,966 in Herpes, 5,099 in Papilloma, 3,044 in Influenza, 1,300 in Hepatitis, 927 in Dengue and 709 in Zika from different public databases (Altunkaya et al., 2017; Ammari et al., 2016; Calderone et al., 2015; Durmuş Tekir et al., 2013; Guirimand et al., 2015), as well as 586 interactions of SARS-CoV-2 (Gordon et al., 2020; Li et al., 2020). As for negative sets, we employed ‘Dissimilarity-Based Negative Sampling’ (Eid et al., 2016; Yang et al., 2020, 2021), and sampled non-interacting sets that were 10 times larger than the corresponding interaction data sets (see Materials and methods). While **Table 1** indicates generally high prediction performance of our deep learning approach, we observed that small sizes of training data sets such as Dengue, Zika and SARS-CoV-2 pointed to decreasing prediction performance.

**Table 1.** Performance of our deep learning architecture (PSSM+CNN+MLP) using 5-fold cross validation.

| Human-virus PPI<br>dataset | Accuracy<br>(%) | Precision<br>(%) | Sensitivity<br>(%) | Specificity<br>(%) | F1-score<br>(%) | AUPRC |
| --- | --- | --- | --- | --- | --- | --- |
| Human-HIV | 98.65 | 95.16 | 89.72 | 99.54 | 92.36 | 0.974 |
| Human-Herpes | 95.26 | 77.11 | 68.10 | 97.98 | 72.33 | 0.768 |
| Human-Papilloma | 95.98 | 82.70 | 70.48 | 98.53 | 76.10 | 0.818 |
| Human-Influenza | 96.10 | 84.22 | 70.30 | 98.68 | 76.63 | 0.834 |
| Human-Hepatitis | 93.43 | 69.27 | 49.77 | 97.79 | 57.92 | 0.636 |
| Human-Dengue | 93.29 | 70.02 | 45.85 | 98.04 | 55.41 | 0.605 |
| Human-Zika | 95.41 | 85.17 | 59.94 | 98.96 | 70.36 | 0.746 |
| Human-SARS-CoV-2 | 90.64 | 45.81 | 16.37 | 98.06 | 24.12 | 0.329 |

As random forest (RF) performs better than other machine learning methods when applied to binary classification problems (Chen et al., 2019; Wu et al., 2009; Yang et al., 2020), we compare the performance of our deep learning approaches (i.e., PSSM+CNN+MLP) to this representative state-of-art classifier. Moreover, we consider three widely-used encoding methods (i.e., LD, CT and AC) for feature representations as the input to the RF classifier using 5-fold cross validation. By comparing AUPRC (area under precision-recall curve) values, we observed that our deep learning method generally outperforms other encoding schemes-based RF classifiers especially when applied to comparatively large datasets (**Table 2**).

**Table 2.** Performance comparison of our deep learning architecture and three sequence encoding scheme-based RF methods using 5-fold cross validation.

| Human-viral PPI dataset | AUPRC |  |  |  |
| --- | --- | --- | --- | --- |
|  | Our method | LD+RF | CT+RF | AC+RF |
| Human-HIV | 0.974 | 0.972 | 0.970 | 0.972 |
| Human-Herpes | 0.768 | 0.741 | 0.737 | 0.699 |
| Human-Papilloma | 0.818 | 0.740 | 0.724 | 0.656 |
| Human-Influenza | 0.834 | 0.813 | 0.795 | 0.713 |
| Human-Hepatitis | 0.636 | 0.571 | 0.580 | 0.537 |
| Human-Dengue | 0.605 | 0.526 | 0.505 | 0.456 |
| Human-Zika | 0.746 | 0.720 | 0.718 | 0.698 |
| Human-SARS-CoV-2 | 0.329 | 0.371 | 0.350 | 0.314 |

To further assess the impact of our encoding scheme to represent the features of interacting proteins, we compared the performance of our deep learning architecture based on PSSM to a different word embedding technique. Specifically, the word2vec encoding method (Chen et al., 2019; Le and Mikolov, 2014) considers each amino acid as a word, and learns a word-embedding of sequences based on a large corpus, where each amino acid is encoded by a 5-dimensional vector. As 20 amino acids are clustered into 7 groups based on their dipoles and volumes of the side chains (Shen et al., 2007; Sun et al., 2017), that were used for the construction of conjoint triads (CT) we one-hot encoded each amino acid of the corresponding sequence. As a result, word2vec+CT one-hot is the concatenation of pre-trained amino acid embeddings where each protein is represented by a *n* × 12 dimensional array. As noted previously, we considered a fixed sequence length of *n* = 2,000 and zero-padded smaller sequences. Training our CNN+MLP approach with word2vec+CT one hot encodings of the corresponding protein sequences, we observed that the representation of sequences through PSSM in our approach provided better prediction performance especially in relatively small datasets such as Dengue, Zika and SARS-CoV-2 (**Table 3**).

**Table 3.**
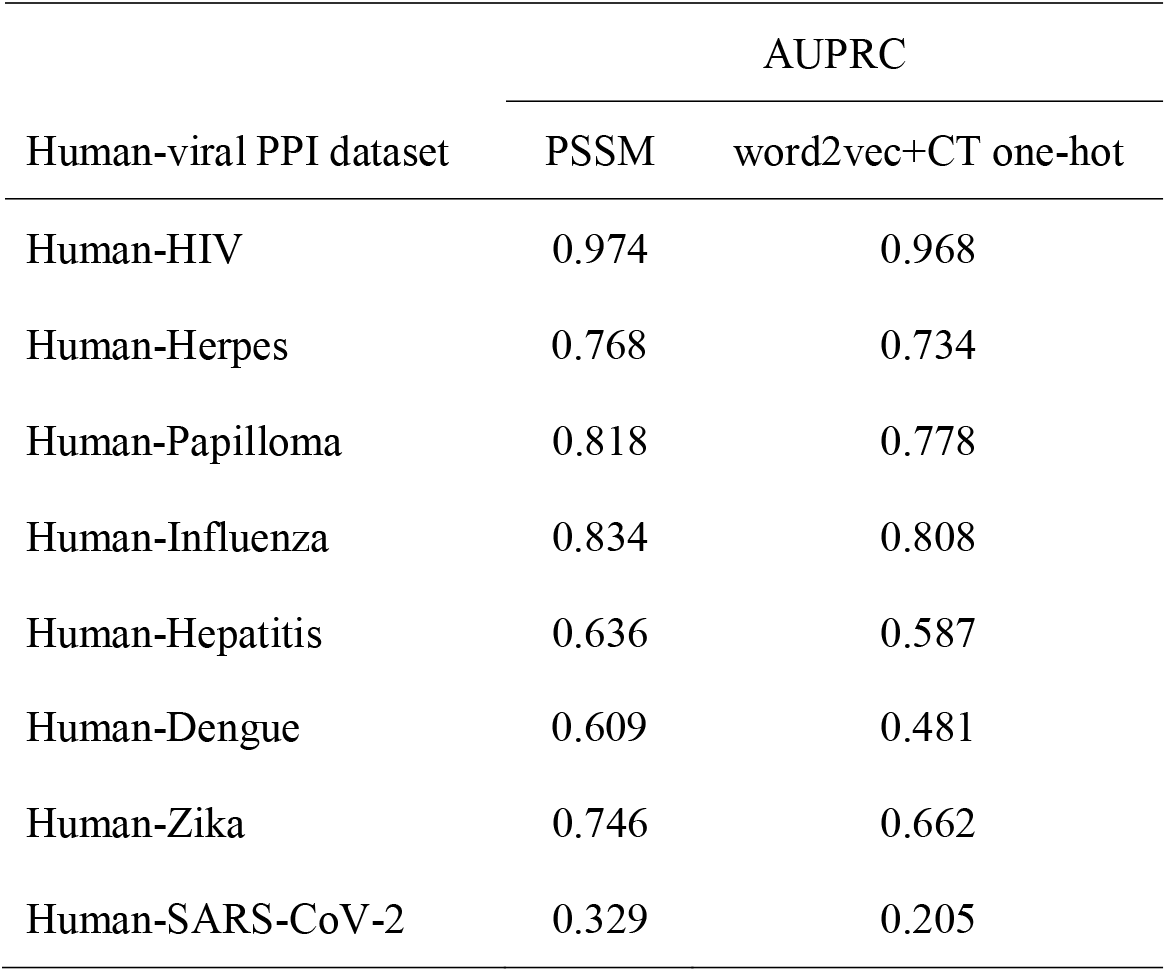
Performance comparison of combinations of different feature encodings (PSSM, word2vec+CT one-hot) and our deep learning algorithm (CNN+MLP).

| Human-viral PPI dataset | AUPRC |  |
| --- | --- | --- |
|  | PSSM | word2vec+CT one-hot |
| Human-HIV | 0.974 | 0.968 |
| Human-Herpes | 0.768 | 0.734 |
| Human-Papilloma | 0.818 | 0.778 |
| Human-Influenza | 0.834 | 0.808 |
| Human-Hepatitis | 0.636 | 0.587 |
| Human-Dengue | 0.609 | 0.481 |
| Human-Zika | 0.746 | 0.662 |
| Human-SARS-CoV-2 | 0.329 | 0.205 |

### Comparison with other existing human-virus PPI prediction methods

To further assess the performance of our proposed method, we compared our method with three existing human-virus PPI prediction approaches. Recently, we proposed a sequence embedding-based RF method to predict human-virus PPIs with promising performance (Yang et al., 2020). In particular, we applied an unsupervised sequence embedding technique (i.e., doc2vec) to represent interacting protein sequences as low-dimensional vectors with rich features that are subjected to RF to predict the presence/absence of an interaction. In Alguwaizani et al.’s work (Alguwaizani et al., 2018), the authors utilized a Support Vector Machine (SVM) model to predict human-virus PPIs based on feature-encoding of protein sequences through patterns of repeats and local amino acid combinations. As for the DeNovo method (Eid et al., 2016), the authors introduced a domain/linear motif-based SVM approach to predict human-virus PPIs. To make a fair comparison, we first constructed the PSSM of the protein sequences of DeNovo’s PPI dataset and used their training set to train our Siamese-based CNN model. Finally, we assessed the performance of our reconstructed deep learning model on the test set provided in (Eid et al., 2016) including 425 positive and 425 negative samples. Furthermore, we evaluated the performance of our previous RF based prediction method and Alguwaizani et al.’s SVM approach with these data sets as well. Notably, **Table 4** suggests that our deep learning and our previously published RF-based method clearly outperformed other approaches.

**Table 4.** Performance comparison of our model (PSSM+CNN+MLP) with existing human-virus PPI prediction methods.

| Method | Accuracy<br>(%) | Precision<br>(%) | Sensitivity<br>(%) | Specificity<br>(%) | F1-score<br>(%) |
| --- | --- | --- | --- | --- | --- |
| Our model | 94.12 | 97.23 | 90.82 | 97.41 | 93.92 |
| Our previous doc2vec+RF<sup>a</sup> | 93.23 | 95.99 | 90.33 | 96.17 | 93.07 |
| Alguwzizani et al.'s SVM<sup>b,d</sup> | 86.47 | 86.56 | 86.35 | 86.59 | NA |
| DeNovo<sup>c,d</sup> | 81.90 | NA | 80.71 | 83.06 | NA |

<sup>a</sup> The corresponding values were retrieved from (Yang et al., 2020). <sup>b</sup> The corresponding values were retrieved from (Alguwaizani et al., 2018). <sup>c</sup> The corresponding values were retrieved from (Eid et al., 2016). <sup>d</sup> NA means the corresponding parameter is not available.

### Cross-viral tests and transfer learning

To explore potential factors that affect prediction performance in a cross-viral setting, we trained our deep learning model on one human-virus PPI data set and predicted protein interactions in a different human-virus system. Utilizing 5-fold cross validation, we expectedly observed that the performance of such naïve cross-viral tests dropped considerably compared to training and testing in the same human-virus system (**Figure 2a**). To allow reliable cross-viral predictions of PPIs, we introduce two transfer learning methods (i.e., ‘frozen’ and ‘fine-tuning’) where we first train the parameters of CNN layers on a source human-virus PPI dataset. Subsequently, we transferred all parameters of CNN layers to initialize a new model (‘frozen’ or ‘fine tuning’) with randomly initialized MLP layers to train on a given target human-virus PPI domain. To comprehensively test our transfer learning approaches, we considered each combination of human-viral PPI sets as source and target domains. **Figure 2b** indicates that a relatively rigid transfer learning methodology by keeping the parameters of the CNN module untouched (i.e., ‘frozen’) and training the MLP layers strongly outperformed the naïve baseline performance as shown in **Figure 2a**. In turn, fine-tuning parameters in the CNN module and training the MLP layers as well with a given target human-viral domain allowed for another increase in performance (**Figure 2c**). As for individual pairs of human-viral domains, we also observed that independently from the training domain the ‘frozen’ transfer methodology worked better compared to the ‘fine-tuning’ approach when the target domain data set was extremely small (i.e., human-SARS-CoV-2). In turn, performance of the ‘frozen’ transfer learning approach dropped compared to ‘fine-tuning’ when the target human-viral domain datasets of PPIs were larger such as human-Hepatitis, human-Dengue and human-Zika.

**Figure 2.**
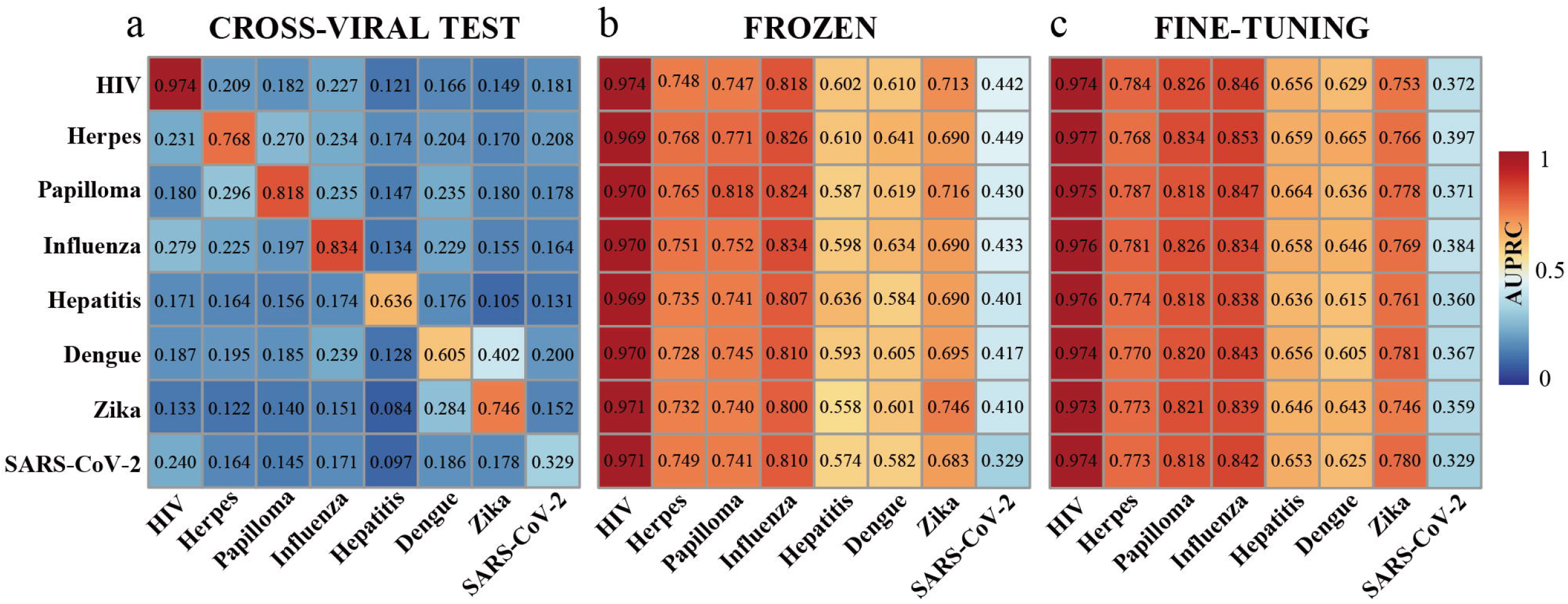
**a** Investigating prediction performance we trained our deep learning model on one human-virus PPI data set (rows) and predicted protein interactions in a different human-virus system (columns). Expectedly, the performance of such cross-viral tests dropped considerably in performance compared to training and testing in the same human-viral system. **b** In a transfer learning step we trained the parameters of the CNN and MLP layers on a source human-virus PPI dataset (rows) and transferred all parameters of CNN layers to initialize a new model to train on a target human-virus PPI dataset (columns). Notably, a relatively rigid transfer learning methodology where we trained the MLP layers with randomly initialized parameters but left the parameters of the feature encoding CNN untouched (i.e., ‘frozen’) strongly outperformed baseline performance in **a**. **c** In turn, fine-tuning parameters where we trained the MLP layers and retrained the CNN layers using a given target human-viral domain allowed for another marked increase in performance.

### Prediction and analysis of human-SARS-CoV-2 PPIs based on transfer learning models

To predict a genome-wide map of potential interactions between proteins of the human host and SARS-CoV-2, we first trained parameters of the CNN layers of our deep learning model utilizing all human-viral protein interactions. Subsequently, we used our two transfer learning approaches to train our set of interactions between proteins of human and SARS-CoV-2. Applying 5-fold cross validations, we observed that the AUPRC of 0.483 with the ‘frozen’ transfer learning approach outperformed the corresponding value of 0.435 when we used the ‘fine-tuning’ method. In addition, training on all source human-virus PPI datasets showed best performance compared to separately training with virus-specific source PPI datasets (data not shown). Therefore, we employed the five ‘frozen’ models of the 5-fold cross-validation based on human-all virus source dataset to predict human-SARS-CoV-2 PPIs and averaged the scores of the five models as the prediction result. At a false positive rate control of 0.01, we identified 946 high-confidence protein interactions between 21 SARS-CoV-2 proteins and 551 human proteins (**Supplementary Table S1**). As for topological network analysis, we found a power-law distribution when we counted the number of human proteins that were targeted by a certain number of viral proteins (**Figure 3a**). Such an observation suggests that the majority of human proteins are targeted by one viral protein, while a minority of human host proteins interacts with many viral proteins, a result that is in line with previous observations (Wuchty et al., 2010). As targeted human proteins usually play important topological roles (i.e., hubs) in human PPI networks, we performed an enrichment analysis of our identified viral targets by using a human specific PPI network. In particular, we collected 365,284 human PPIs from the HIPPIE database (Alanis-Lobato et al., 2017) and observed that targeted human proteins are enriched in bins of increasing degree (**Figure 3b**), a result that is consistent with previous findings as well (Dyer et al., 2008; Wuchty et al., 2010). Considering 2,916 human protein complexes from the CORUM database (Giurgiu et al., 2019) we found that viral targets are enriched in sets of proteins that participate in an increasing number of protein complexes (**Figure 3c**) (Wuchty et al., 2010). To illustrate viral similarities, we compared the experimentally known human-SARS-CoV-2 interactome and our predicted interactome with their counterparts of seven other viruses (**Figure 3d**). While our predictions show similar overlaps of viral targets with the experimentally obtained interactomes, we further found that Dengue virus and Influenza virus had the most similar interacting partners in both predictions and experimentally known interactions (*p*-value ≤ 0.05, hypergeometric test). Notably, the association with Influenza virus is of particular interest as this virus also induces respiratory disease (i.e., pneumonia).

**Figure 3.**
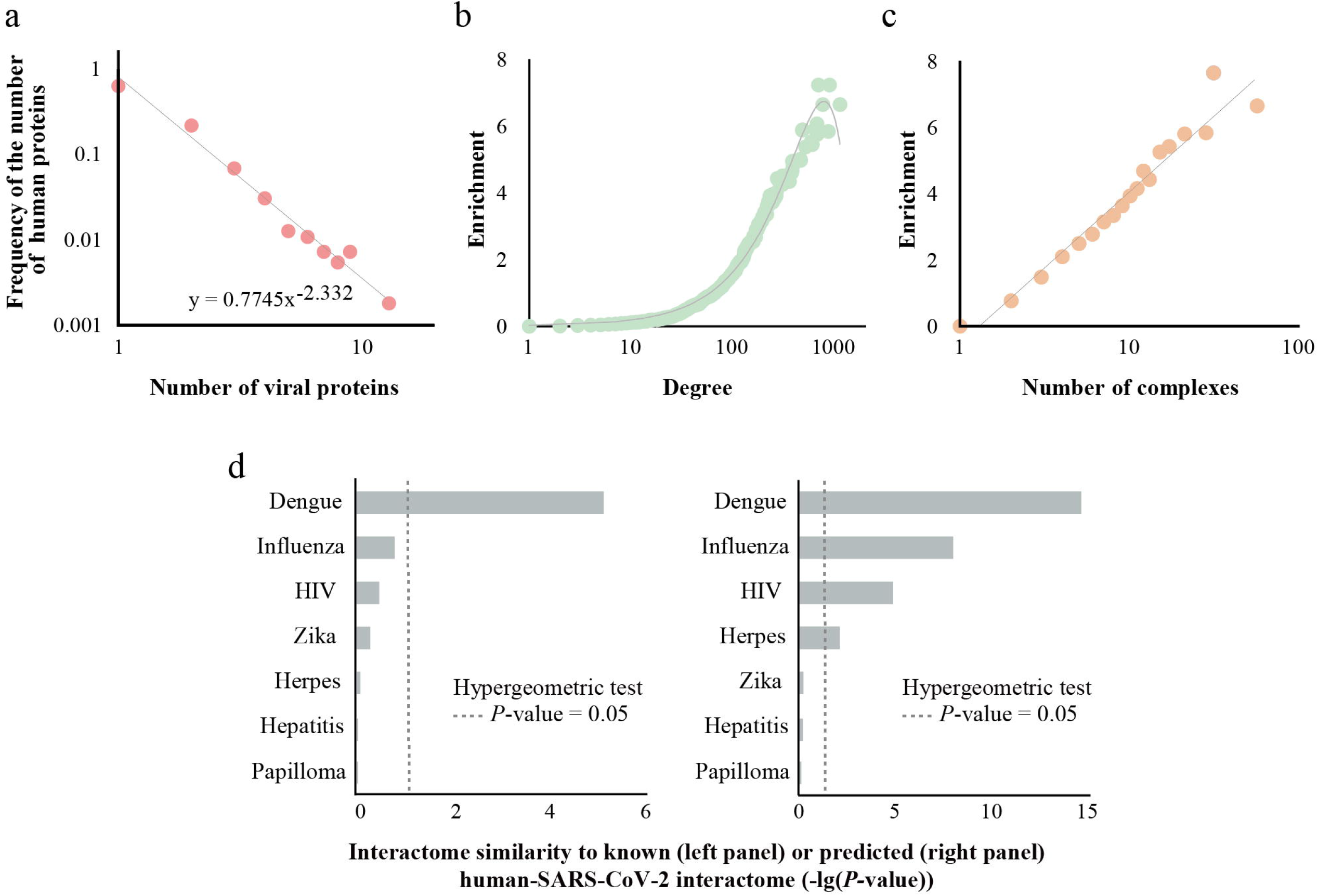
**a** Analyzing the predicted network of human-SARS-CoV-2 protein interactions we observed a power-law in the frequency distribution of the number of human proteins that are targeted by a certain number of viral proteins. **b** Viral targets were enriched in bins of human host proteins that had an increasing number of interactions. **c** Viral targets were enriched in sets of proteins that participate in an increasing number of human protein complexes. **d** Analysis of the significance of the overlap between known (left panel) and predicted (right panel) human interacting proteins of SARS-CoV-2 and known human interacting proteins of other pathogens with a hypergeometric test revealed strong similarities with Dengue, Influenza and HIV. The background gene set for the test consisted of all unique human interacting proteins across all pathogens (N=6,218 and N=6,200 proteins for comparing with known and predicted human interacting proteins of SARS-CoV-2, respectively).

### General functional roles of SARS-CoV-2 targets

On a quantitative functional level, we calculated overlaps between both our predicted and known viral targets and several functional gene sets [i.e., essential gene set, transcription factor set, kinases set and posttranslational modification (PTM) set] to characterize the functions of viral targets (**Figure 4**). As a result, we found that both predicted and known viral targets were significantly enriched with essential genes (**Figure 4a**), while they rarely were transcription factors and kinases (**Figure 4b, c**). In turn, we observed that ubiquitinated, methylated and acetylated proteins increasingly appeared in sets of viral targets while phosphorylated target appeared not enriched (**Figure 4d-g**). In particular, several studies have revealed SARS-CoV-2 targets human proteins that receive PTMs relate to pathways that modulate host antiviral immune responses (Pruimboom, 2020; Shin et al., 2020). In turn, phosphorylation appears ubiquitous and not specific for viral targets.

**Figure 4.**
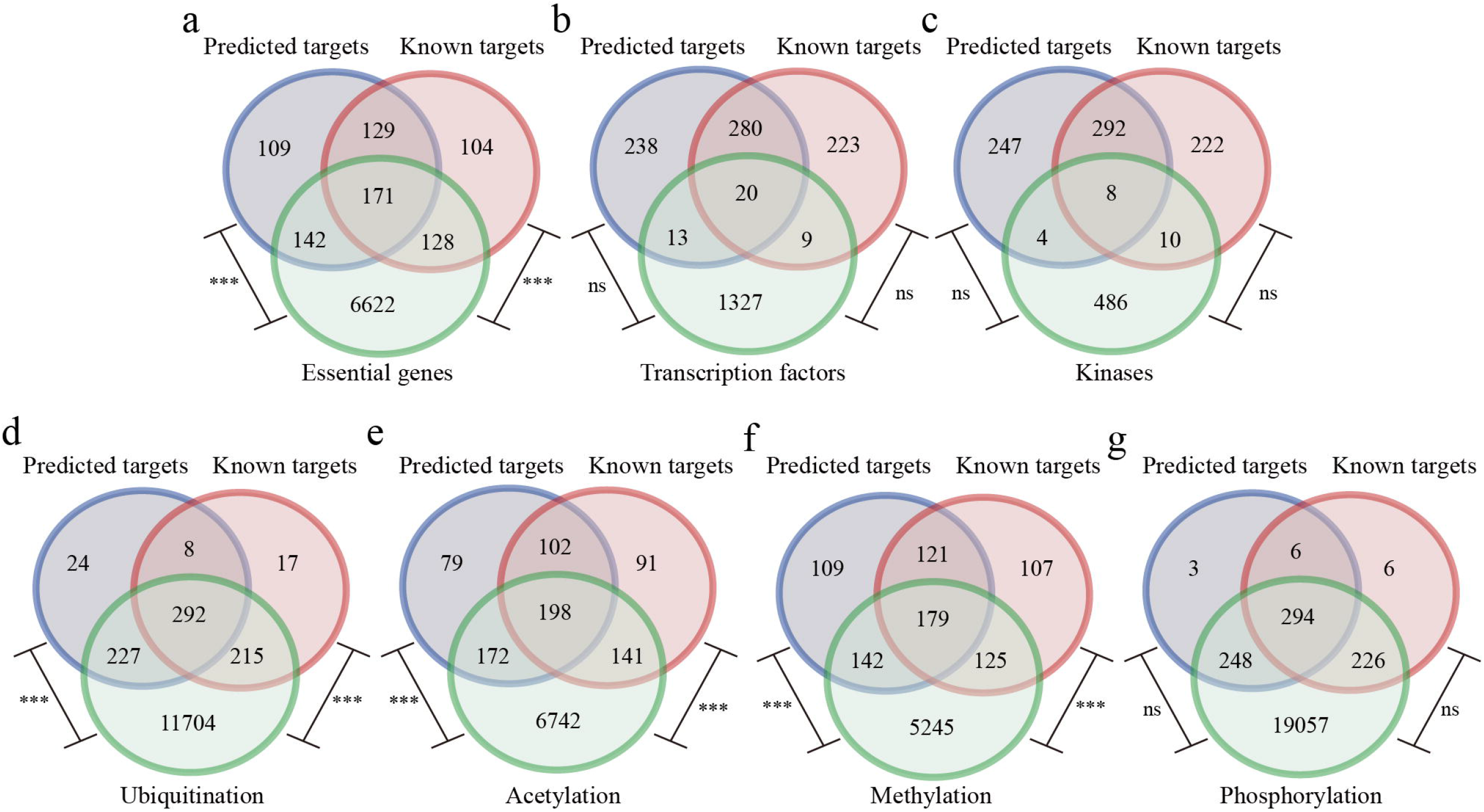
**a** Venn diagram indicates that overlaps of essential genes and known/predicted viral were statistically significant (*** *P*-value ≤ 0.001, hypergeometric test). **b, c** In turn, overlaps of transcription factors/kinases and predicted/known viral targets were not significant (ns *P*-value ≥ 0.05). **d-f** Considering overlaps with post-translational modifications known/predicted viral targets were significantly enriched with ubiquitinated acetylated, and methylated proteins while phosphorylated proteins were not enriched (*** *P*-value ≤ 0.001, ns *P*-value > 0.05).

### Comparative analysis of known and our predicted human-SARS-CoV-2 PPIs

Comparing our predicted and experimentally obtained sets of interactions between proteins of SARS-CoV-2 and the human host, we found considerable overlaps. In particular, 298 out of 946 predicted PPI were identified through previous experimental efforts that amount to 52.5% of known interactions in SARS-CoV-2, while 648 were specifically identified through our deep learning approach (**Figure 5a, Supplementary Table S1**), indicating the reliability and specificity of our model for the identification of novel interactions. To further excavate functional similarities and differences between known and predicted targets, we performed functional and pathway enrichment for experimentally known and predicted viral targets, respectively. Considering hypergeometric tests (Bonferroni corrected *P*-value ≤ 0.01), we observed a relatively large number of shared GO enrichment terms/KEGG enrichment pathways of experimentally confirmed targets and predicted targets, indicating functionally similar between them, which further demonstrates the reliability and quality of our predictions. (**Figure 5b, Supplementary Table S2, S3**). In more detail, enriched GO BP terms in human host proteins that were found in the experimental PPIs and predictions are displayed in **Figure 5c**. Indicating high confidence of our predicted targets to discover functions SARS-CoV-2 meddles in, we found that functional enrichments of experimental and predicted viral targets both point to the involvement of viral targets in protein transport, protein import and mRNA export from the nucleus. Notably, our predictions augment such functions, indicating that the virus may also interfere with nuclear pore organization and assembly as well as protein export from the nucleus.

**Figure 5.**
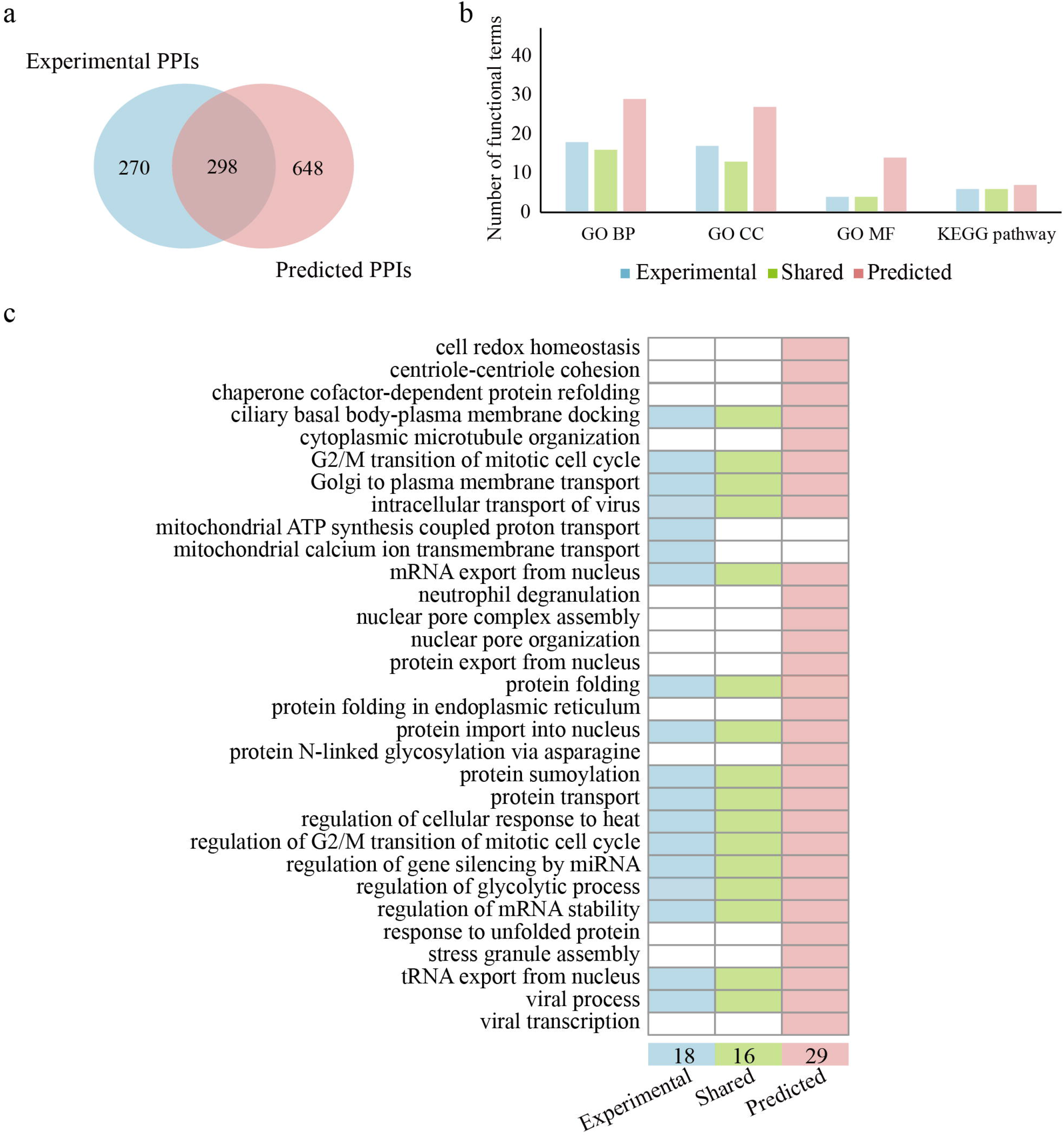
**a** The Venn diagram indicates a sizeable overlap of experimentally observed and predicted interactions between proteins of the human host and SARS-CoV-2. **b** In a quantitative functional analysis of targeted human host proteins, we considered the enrichment of GO terms and KEGG pathways through a hypergeometric test (Bonferroni corrected *P*-value ≤ 0.01). We found that a relatively large shared GO enrichment terms and KEGG enrichment pathways in groups of host proteins that appeared in the experimentally known PPIs and predictions. **c** In more detail, we observed that enriched GO BP terms in host proteins that were found in the experimental PPIs and predictions were functionally similar.

### Modular analysis of human-SARS-CoV-2 PPI network

To further explore potential functional modules that can reveal SARS-CoV-2 biology, we combined our predicted 946 human-SARS-CoV-2 PPIs with known human-specific PPIs as of the HIPPIE database (Alanis-Lobato et al., 2017) (**Figure 6a**). Specifically, we identified 9 topological modules based on connectivity among human proteins (**Figure 6a, b**), utilizing the MCODE algorithm (Bader and Hogue, 2003). Investigating the enrichment of GO BP terms and KEGG pathways through hypergeometric tests (Bonferroni adjusted *p*-value ≤ 0.05), we observed that these modules largely revolve around ribosome biogenesis, retrograde protein transport, elastic fiber assembly, mitochondrial translation, protein processing in endoplasmic reticulum, stress granule regulation, protein folding in endoplasmic reticulum, centrosome and gene splicing (**Supplementary Table S4**).

**Figure 6.**
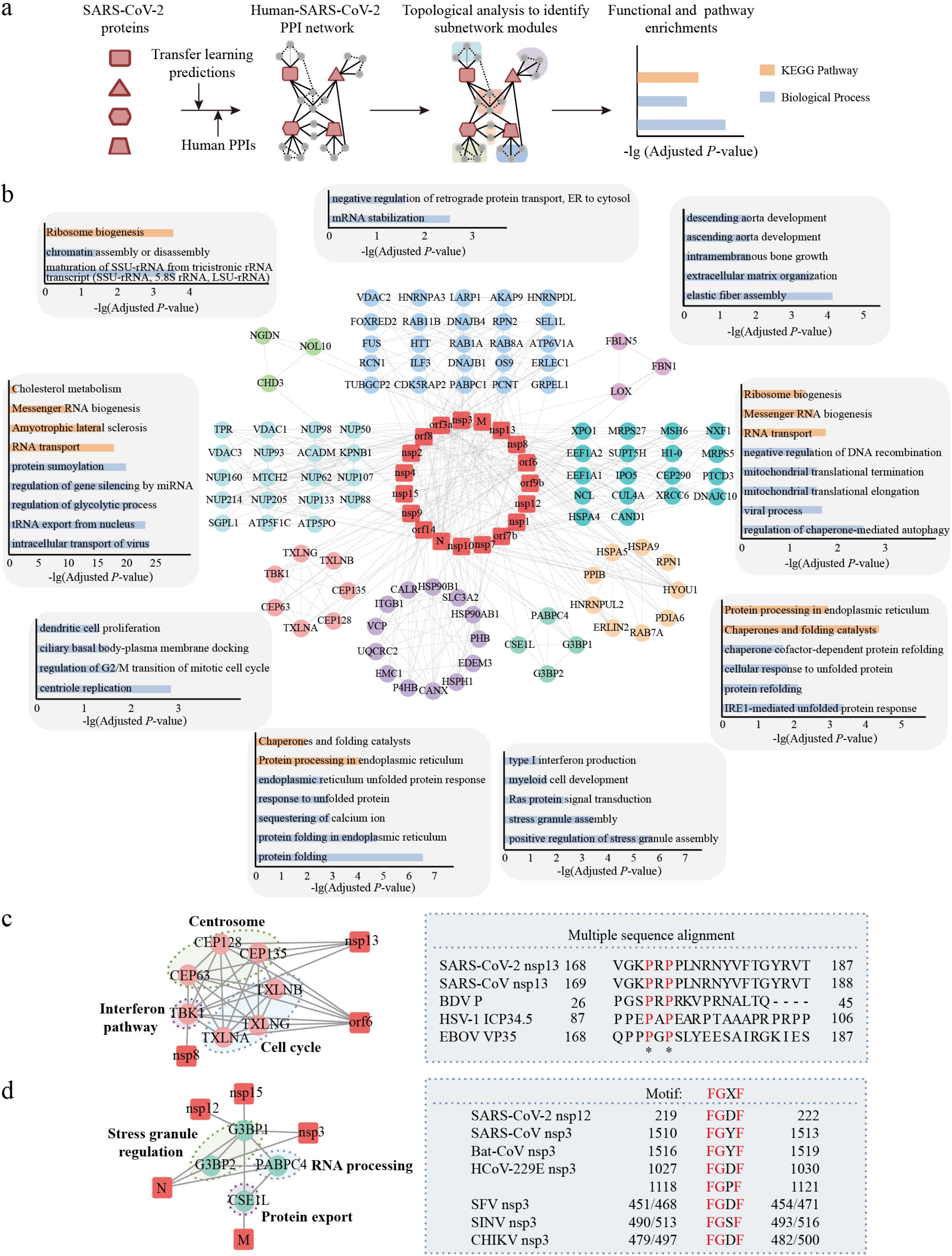
**a** Combining predictions that we obtained with the transfer learning approach and known human PPIs we determined connectivity-based modules that were subjected to functional interpretation. **b** Human-SARS-CoV-2 PPI network with enriched GO BP terms and KEGG pathways for each topological module. **c** SARS-CoV-2 targets a module that involves the centrosome, cell cycle and interferon pathway. SARS-CoV-2 interacts with interferon pathway and presents a conserved region with multiple viral pathogens. A conserved binding motif that nsp13 of SARS-CoV-2 and proteins of various other viral pathogens share suggests that SARS-CoV-2 nsp13 protein may interfere with the regulation processes of IFN, supporting antiviral innate immune response. **d** SARS-CoV-2 targets a module that involves stress granule regulation, RNA processing and protein export, and interacts with stress granule proteins and shows potential interaction patterns by a conserved amino-acid motif in the nsp12 of SARS-CoV-2 and nsp3 proteins of other viruses.

Considering a module that was enriched with centrosome functions through interactions with nsp13 and cell cycle functions through interactions with orf6, we also found that this module harbors human genes that allow SARS-CoV-2 to interact with innate immune pathways which is consistent with previous findings (Gordon et al., 2020). As shown in the module, the interferon (IFN) pathway is targeted though TBK1 by nsp8, nsp13 and orf6, a serine/threonine-protein kinase that plays an important role in the induction of the antiviral IFN responses to foreign agents such as viruses. A number of viral proteins binds to TBK1 and regulates its kinase activity to reduce TBK1-mediated secretion of IFN and induction of an antiviral state, such as Borna disease virus (BDV) P protein (Unterstab et al., 2005), Human herpesvirus 1 (HSV-1) ICTP34.5 protein (Manivanh et al., 2017) and Ebola virus (EBOV) VP35 protein (Prins et al., 2009). BDV P protein itself is phosphorylated by TBK1, suggesting that P functions as a viral decoy substrate that prevents activation of cellular target proteins of TBK1. Furthermore, residues from 87 to 106 in HSV-1 ICTP34.5 protein interact with TBK1 to modulate type I IFN signaling (Manivanh et al., 2017; Verpooten et al., 2009). Considering the multiple sequence alignment of these viral proteins and nsp13 of SARS-CoV and SARS-CoV-2 we found a conserved binding motif (**Figure 6c**), corroborating our assumption that SARS-CoV-2 nsp13 protein may also interfere with the regulation processes of IFN that support antiviral innate immune response.

In a different module (**Figure 6d**), various virus proteins of SARS-CoV-2 target stress granule regulation (N, nsp3, nsp12, nsp15), RNA processing (N) and protein export functions (M). Notably, G3BP1 and G3BP2 are stress granule proteins with anti-viral activity. The nsp3 protein of several other viral pathogens interacts with G3BP1 via C-terminus including Sindbis virus (SINV), Semliki forest virus (SFV) and Chikungunya virus (CHIKV). Such interactions inhibit the formation of host stress granules on viral mRNAs while the nsp3-G3BP1 complexes bind viral RNAs and probably orchestrate the assembly of viral replication complexes. In particular, coronavirus commonly manipulates the stress granules and related RNA biology processes while stress granule formation/assembly is considered a primary antiviral response (Nakagawa et al., 2018; Quispe-Tintaya, 2019). To corroborate such findings, we found a conserved amino-acid motif (‘FGXF’) in the nsp3 proteins of these viruses. In particular, mutations in the ‘FGDF’ motif of SFV nsp3 protein indicated that residues F, G, F are essential for G3BP-binding (Panas et al., 2015). Notably, we found that the SARS-CoV-2 nsp12 protein and the SARS-CoV nsp3 protein both contained the complete ‘FGXF’ motif, indicating the reliability of our prediction and the detailed interaction pattern.

## Discussion and conclusions

We designed a Siamese-based multi-scale CNN architecture by using PSSM to represent the sequences of interacting proteins, allowing us to predict interactions between human and viral proteins with an MLP approach. In comparison, we observed that our model outperformed previous state-of-the-art human-virus PPI prediction methods. Furthermore, we confirmed that the performance of the combination of our deep learning framework and the representation of the protein features as PSSM was mostly superior to combinations of other machine learning and pre-trained feature embeddings. While we found that our model that was trained on a given source human-viral interaction data set performed dismally in predicting protein interactions of proteins in a target human-virus domain in a naïve way, we introduced two transfer learning methods (i.e., ‘frozen’ type and ‘fine-tuning’ type). Such methods allowed us to train on a source human-virus domain and retrain the layers of CNN with data of a target domain. Notably, our methods increased the cross-viral prediction performance dramatically, compared to the naïve baseline model. In particular, for small target datasets, fine-tuning pre-trained parameters that were obtained from larger source sets increased prediction performance. Specifically, keeping the CNN layers from large source datasets untouched improved the generalization ability of the new model (i.e., ‘frozen’ model), while retraining layers of CNN (i.e., ‘fine-tuning’ model) allowed the specificity of predictions in target domain datasets.

Finally, we employed our ‘frozen’ transfer learning to predict human-SARS-CoV-2 PPIs and performed in-depth network analysis based on the identified interactions. Our transfer learning model resembled closely the functions and characteristics of experimentally obtained interactions and indicated novel functions that the virus potentially targets. Furthermore, we identified several high-confidence interactions involved in various vital immune pathways and anti-viral processes. Taken together, our transfer learning method can be effectively applied to predict human-virus PPIs in a cross-viral setting and the study of viral infection mechanism.

## Materials and methods

### Deep learning network architecture

#### Pre-acquired protein sequence profile module

For each protein sequence with variable lengths, we generated a sequence profile, called position specific scoring matrix (PSSM). In particular, we performed PSI-BLAST searches with default parameters applying a threshold of E-value < 0.001 in the UniRef50 protein sequence database (Suzek et al., 2015) as PSI-BLAST allows us to discover protein sequences that are evolutionarily linked to the search sequence (Hamp and Rost, 2015; Hashemifar et al., 2018). Sequence profiles thus obtained for each search sequence were processed by truncating profiles of long sequences to a fixed length *n* and zero-padding short sequences, a method widely used for data pre-processing and effective training (Matching, 2018; Min et al., 2017). As a result, we obtained a *n* × 20 dimensional array *S* for each protein,

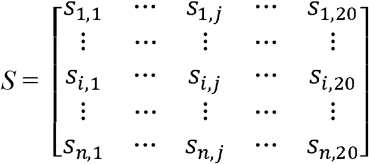

capturing the probability *s_i,j_* that the residue in the *i*^th^ position of the sequence is the *J^th^* out of the alphabet of 20 amino acids.

#### Siamese CNN module

To capture complex relationship between two proteins we employ a Siamese CNN architecture with two identical CNN sub-networks that share the same parameters for a given pair of protein profiles *S, S*’. Each sub-network produces a sequence embedding of a single protein profile that are then concatenated. While each single CNN module consists of a convolutional and pooling layer, we leveraged four connected convolutional modules to capture the patterns in an input sequence profile.

Specifically, we use *X*, a *n*× *s* array of length *n* with *s* features in each position. The convolution layer applies a sliding window of length *w* (the size of filters/kernels) to convert *X* into a (*n* — *w* + 1) × *fn* array *C* where *fn* represents the number of filters/kernels. *C_i,k_* denotes the score of filter/kernel *k*, 1 ≤ *k* ≤ *fn*, that corresponds to position *i* of array *X*. Moreover, the convolutional layer applies a parameter-sharing kernel *M*, a *fn* ×*m* × *s* array where *M_k,j,l_* is the coefficient of pattern *k* at position *j* and feature *l*. The calculation of *C* is defined as

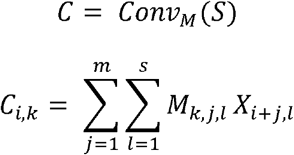

Furthermore, the pooling layer is utilized to reduce the dimension of *C* to a (*n* — *p* + 1) × *fn* array *P* where *p* is the size of pooling window. Array *P* = *Pool*(*C*) is calculated as the maximum of all positions *i* < *j* ≤ *i* + *p* over each feature *k* where 1 ≤ *i* ≤ (*n* — *m* + 1) — *p*,

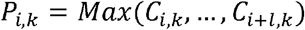

#### Prediction module

The prediction module concatenates a pair of protein sequence embedding vectors into a sequence pair vector as the input of fully connected layers in an MLP and computes the probability that two proteins interact. The MLP contains three dense layers with leaky ReLU where cross-entropy loss is optimized for the binary classification objective defined as

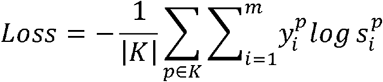

where *y_i_* is numerical class label of the protein pair *p*. The output of the MLP for the protein pair *p* is a probability vector 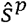, whose dimensionality is the number of classes *m. s* is normalized by a softmax function, where the normalized probability value for the *i^th^* class is defined as 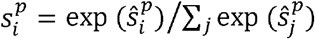.

#### Implementation details

As for pre-acquired sequence profile construction, we consider a fixed sequence length of 2,000. As for the construction of our learning approach, we employ four convolutional modules, with input sizes 20, 64, 128 and 256. The convolution kernel size is set to 3 while the size of pooling window is set to 2 with 3 max-pooling layers and a global max-pooling layer. To optimize the cross-entropy loss function we used AMSGrad (Reddi et al., 2018) and set the learning rate to 0.0001. The batch size was set to 64, while the number of epochs was 100. The fully connected layers contained three dense layers with input size 1,024, 512, 256 and output a two-dimensional vector with the last softmax layer. The whole procedure was implemented with keras (https://keras.io/) with GPU configuration.

### Data set construction

We collected experimentally verified human-virus PPI data capturing 31,381 interactions in all viruses. Specifically, we accounted for 9,880 interactions in HIV, 5,966 in Herpes, 5,099 in Papilloma, 3,044 in Influenza, 1,300 in Hepatitis, 927 in Dengue and 709 in Zika from five public databases, including HPIDB (Ammari et al., 2016), VirHostNet (Guirimand et al., 2015), VirusMentha (Calderone et al., 2015), PHISTO (Durmuş Tekir et al., 2013) and PDB (Altunkaya et al., 2017). As for interactions of proteins of SARS-CoV-2, we used two recently published interaction sets (Gordon et al., 2020; Li et al., 2020) that captured 291 and 295 PPIs, respectively. To obtain high-quality PPIs, we removed interactions from large-scale MS experiments that were detected only once, non-physical interactions and interactions between proteins without available PSSM features.

Sampling negative interactions, we employed a ‘Dissimilarity-Based Negative Sampling’ method (Eid et al., 2016) as outlined in our previous work (Yang et al., 2021, 2020) and set the ratio of pos./neg. samples to 1:10. The key strategy of the ‘negative sampling’ is if the sequences of viral protein A and B are similar, and A interacts with human protein C (i.e., A-C is a positive sample), B-C is not selected as a negative sample.

### Two types of transfer learning methods

To further improve the performance of our deep neural network especially when dealing with smaller datasets, we propose two transfer learning methods that keep the parameters of the CNN layers constant (i.e., ‘frozen’) or allow their fine-tuning in the early layers (i.e., ‘fine-tuning’) and applied them to eight human-virus PPI sets. In more detail, we used the proposed DNN architecture to train the models based on a given source set of human-virus interactions to obtain pre-trained parameters in the CNN layers that learn the representation of the protein sequences. In subsequent transfer learning steps, we keep the parameters of these CNN layers constant (i.e., frozen) and only train parameters of the fully connected layers of the MLP to predict interactions in a target human-viral interaction set. As an alternative, our ‘fine-tuning’ approach trains parameters of the fully connected layers of the MLP and retrains the parameters of CNN layers that we obtained from the initial training step and change such parameters by learning the interactions in a target set of human-virus interactions. The detailed parameter selection and optimization is shown in **Supplementary Table S5**.

### Alternative machine learning and feature encoding methods

#### Random Forest

RF (Hamp and Rost, 2015; Wu et al., 2009) is an ensemble learning method where each decision tree is constructed using a different bootstrap sample of the data (‘bagging’). In addition, RF changes how decision trees are constructed by splitting each node, using the best among a subset of predictors randomly chosen at that node (‘boosting’). Compared to many other classifiers this strategy turns out to be robust against over-fitting, capturing aggregate effects between predictor variables. We utilize the GridSearchCV function to optimize the parameters for the RF algorithm and set the ‘neg_log_loss’ scoring function as the assessment criterion. The detailed parameter selection and optimization is shown in **Supplementary Table S5**.

#### Alternative feature encoding approaches

Amino acid sequences provide primary structure information of a protein that work well as feature representations of binary PPIs. Here, we use three commonly used sequence-based encoding schemes including Local Descriptors (LD) (Davies et al., 2008; Yang et al., 2010), Conjoint Triads (CT) (Sun et al., 2017), and Autocorrelation (AC) (Guo et al., 2008; You et al., 2013). Generally, these features cover specific, yet different aspects of protein sequences such as physicochemical properties of amino acids, frequency information of local patterns, and positional distribution information of amino acids.

### Evaluation criteria

We conduct 5-fold cross-validation to evaluate the performance of predictive models. Under 5-fold cross validation, all the PPI datasets are equally divided into five non-overlapping subsets and each subset owns once chance to train/test the model which can provide an unbiased evaluation. We aggregate the following metrics [i.e., accuracy, precision, sensitivity, specificity, F1-score, and area under the precision recall curve (AUPRC)] to evaluate the performance of the proposed method. In particular, we define accuracy as 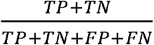, precision as 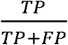, sensitivity/recall as 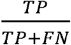 specificity as 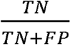 and the F1-score as 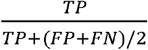, where TP/FP is the number of true/false positives and TN/FN is the number of true/false negatives.

### Enrichment analysis

#### Enrichment analysis of viral targets in degree and human complexes

As for degree of human proteins, we obtained 365,284 human PPIs from HIPPIE database (Alanis-Lobato et al., 2017) and calculated the degree of each viral target based on the compiled PPI network. Following the enrichment approach in (Devkota and Wuchty, 2020), the enrichment of viral targets was determined as a function of degree *k*. In particular, *N_≥k_* is the number of viral targets with degree ≥ *k.* Randomly sampling *N_≥k_* proteins from the whole set of viral targets, we recalculated the corresponding random number of proteins with degree ≥ *k* (i.e., 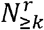). Note, that the random sampling was conducted 100 times and an average of 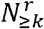 was thus obtained. The enrichment of viral targets that own at least degree *k* was then defined as 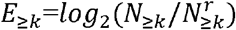 In general, *E_≥k_* > 0 points to an enrichment of viral targets with degree ≥ *k*.

Regarding human complexes, we collected 2,916 human complexes from CORUM (Giurgiu et al., 2019) and calculated the number of participated human complexes of each viral target. Analogously, the enrichment of viral targets as a function of the number of participated human complexes was determined.

#### GO enrichment analysis

GO annotation data of human proteins were downloaded from http://current.geneontology.org/ (The Gene Ontology Consortium, 2019). Using all human proteins mapped to three categories of GO terms [i.e., Cellular Component (CC), Biological Process (BP) and Molecular Function (MF)] as reference sets, enriched GO terms of viral targets were determined by hypergeometric tests, where corresponding *P*-values were Bonferroni corrected.

#### KEGG pathway enrichment analysis

KEGG pathway data were downloaded from https://www.genome.jp/kegg/ (Kanehisa and Goto, 2000). Using all human proteins in KEGG pathways as a reference set, enriched KEGG pathways of viral targets were identified by hypergeometric tests where corresponding *P*-values were Bonferroni corrected.

#### Essential genes, transcription factors, kinases and post-translational modifications (PTM)

We collected 7,063 human essential genes from the Online GEne Essentiality database (OGEE) (Chen et al., 2012) and the Database of Essential Genes (DEG) (Luo et al., 2020). We obtained 1,369 human transcription factors from Gene Transcription Regulation Database (GTRD) (Yevshin et al., 2019) and 508 human kinases from Kinome NetworkX database (Cheng et al., 2014). As for PTM information we obtained 11,670 ubiquitinated proteins, 17,351 phosphorylated proteins, 5,420 methylated proteins and 6,992 acetylated proteins from PhosphoSitePlus database (Hornbeck et al., 2015). As for the enrichment analysis, we employed hypergeometric tests for each PTM type while all 20,251 reviewed human proteins as of the SwissProt database are considered as the reference set.

### Module identification and functional analysis of the modules

In order to build the integrated interaction network for topological analysis, we first collected known protein interactions between the human proteins predicted to interact with SARS-CoV-2 from the HIPPIE database (Alanis-Lobato et al., 2017). In the next step, we combined our predicted human-SARS-CoV-2 PPIs with the known human PPIs into the final interaction network. Moreover, we clustered human proteins within the network based solely on their topological connectivity. We applied the MCODE plugin (Bader and Hogue, 2003) to find clusters of densely interconnected human proteins denoting potential functional modules or parts of pathways (include loops: no; degree cutoff: 2; haircut: no; fluff: no; node score: 0.4; kcore: 2; max depth: 100). Visualizations of the modules (i.e., subnetworks) were carried out with Cytoscape (Shannon et al., 2003). Enrichment analysis for each cluster was performed by using hypergeometric tests, where corresponding *P*-values were Bonferroni corrected, and only the five most enriched GO BP terms and KEGG pathways were considered (Adjusted *P*-value ≤ 0.05) in Figure 6. All the enriched GO terms and KEGG pathways are listed in Supplementary Table S4.

## Supplementary data

**Supplementary Table S1.** Experimentally verified and predicted human-SARS-CoV-2 PPIs.

**Supplementary Table S2.** Enriched GO terms for experimental and predicted SARS-CoV-2 interacting proteins.

**Supplementary Table S3** Enriched KEGG pathways for experimental and predicted SARS-CoV-2 interacting proteins.

**Supplementary Table S4** Enriched GO terms for identified topological modules.

**Supplementary Table S5** Parameter selection and optimization of different frameworks.

## Acknowledgements

We are grateful to those scientists/developers who enabled the development of the deep learning method by making their data/databases/software freely accessible to the community.

## Availability of data and materials

Source code and datasets are available in https://github.com/XiaodiYangCAU/TransPPI/.

## Authors’ Contributions

SW, XY and ZZ conceived the study and designed the experiment. XY performed the computational analysis and wrote the manuscript. SW and ZZ supervised the study and revised the manuscript. XL, SY and SW provided data preparation and analysis. All authors have read and approved the final manuscript.

